# A convergent structure-function substrate of cognitive imbalances in autism

**DOI:** 10.1101/2021.01.14.426284

**Authors:** Seok-Jun Hong, Laurent Mottron, Bo-yong Park, Oualid Benkarim, Sofie L. Valk, Casey Paquola, Sara Larivière, Reinder Vos de Wael, Janie Degré-Pelletier, Isabelle Soulieres, Bruce Ramphal, Amy Margolis, Michael Milham, Adriana Di Martino, Boris C. Bernhardt

## Abstract

Autism is a common neurodevelopmental condition characterized by substantial phenotypic heterogeneity, which hinders diagnosis, research, and intervention. A leading example can be found in marked imbalances in language and perceptual skills, where deficits in one domain often co-exist with normal or even superior performance in the other domain. The current work capitalized on multiple data analytics including data-driven subtyping and dimensional approaches to quantify cognitive imbalances in multi-site datasets of individuals diagnosed with autism spectrum disorder (ASD) and neurotypical controls, and assessed structural and functional brain network substrates. Studying cognitive dimensions as well as multimodal neuroimaging signatures in 155 ASD and 151 neurotypical individuals, we observed robust evidence for a structure-function substrate of cognitive imbalances in ASD. Specifically, ASD presented with marked imbalances in cognitive profiles relative to neurotypical controls, characterized by verbal to non-verbal intelligence discrepancy. Different analytical approaches including subtyping and dimensional regression methods converged in showing that these imbalances were reflected in atypical cortical thickening and functional integration of language networks, alongside with sensory and higher cognitive networks. Phenotypic findings could be replicated in an independent sample of 325 ASD and 569 neurotypical controls. Although verbal and non-verbal intelligence are currently considered as specifiers unrelated to the categorical diagnosis of autism, our results show that intelligence disparities are accentuated in ASD and relate to a consistent structure-function substrate affecting multiple brain networks. Our findings motivate the incorporation of cognitive imbalances in future autism research, which may help to parse the phenotypic heterogeneity of autism and potentially inform intervention-oriented subtyping.

## Introduction

Autism spectrum disorder (ASD) is a prevalent neurodevelopmental condition currently diagnosed in more than 1 in 54 children (1). A mounting literature suggests that one of the most critical challenges to diagnostic and research advances in autism is phenotypic heterogeneity in various cognitive abilities and sensory modalities (2–4). It ranges from deficits in adaptive, language, or perceptual functions to normal, or even enhanced abilities compared to typically developing individuals, suggesting that the ‘disorder’ term inherent to ASD may not equally qualify for all diagnosed individuals (5,6). Merging all individuals on the spectrum can, thus, miss clinically important subgroups of ASD, and potentially impart null (7) or at best conflicting neuroimaging and cognitive findings (3,8). Ultimately, these challenges hinder our ability to understand functional impairments, underlying mechanisms, and the development of more effective therapy.

The heterogeneity of ASD is particularly apparent in perceptual (or non-verbal) and verbal skills (9) across affected individuals. Prior studies show that the discrepancy between verbal and non-verbal intelligence quotient (IQ) measures is more prevalent among children with ASD than those with typically development (10–13), suggesting that IQ imbalance may be an important characteristic of ASD. Such differences seem plausible, also given that individuals with ASD who experience speech onset delay and are minimally verbal may perform adequately, or even better than neurotypicals, on tasks that do not require verbal skills and target non-verbal perceptual reasoning (5). Conversely, individuals with ASD who present with normal or reduced sensory-perceptual processing may show verbal abilities that are comparable to neurotypicals, or at higher levels (14). Imbalances between verbal and nonverbal skills has been suggested to reflect atypical developmental pathways in ASD, and prior studies have begun to assess cognitive imbalances in ASD by studying the ratio of verbal to nonverbal IQ (vnIQ) measures, which overall suggest that IQ discrepancies may represent a common alteration in ASD (10,15).

While neural substrates underlying the cognitive imbalances in ASD remain to be investigated, it has been postulated that different functional systems may atypically compete during brain development in individuals with ASD (16). Such a broad mechanism may simultaneously affect the organization of multiple cortical areas, including sensory-motor, language, as well as higher cognitive regions that would otherwise show typical areal specialization in normal brain development (17). Neuroimaging, notably multimodal magnetic resonance imaging (MRI), allows for the study of typical and atypical brain organization and development *in vivo* (18–21). In particular, surface-based quantitative MRI analysis can be used to assess morphological variations of cortical areas (22–25), and resting-state functional MRI (rs-fMRI) analysis can probe cortico-cortical connectivity (26–30). In ASD, prior neuroimaging studies have explored morphological and connectome abnormalities. At the level of brain structure, several recent studies in multi-site samples have converged on patterns of cortical thickening in ASD, in a spatial distribution affecting largely frontal and temporal lobe regions (23,25,31,32). Findings have nevertheless been somewhat inconsistent, with other work showing cortical thinning (33), no findings (34), or findings of distinct neuroanatomical patterns across different ASD subgroups ranging from cortical thickening to thinning (22). Heterogeneous patterns have also been reported at the level of functional connectivity. Indeed, prior rs-fMRI studies in ASD have suggested a mosaic of connectivity anomalies, with both under-as well as over-connectivity in ASD groups relative to neurotypicals (26,30). While there are studies overall giving an optimistic note on the consistency of the spatial distribution of connectivity anomalies (35,36), other work has pointed to variability in findings that may stem from methodological vibrations as well as sample-specific inclusion criteria (30,37,38).

Despite prior neuroimaging work thus pointing to atypical structure and functional connectivity in ASD overall, studies rarely took into consideration cognitive discrepancies as potential modulating factors of brain organization in this condition. Notably, work in healthy individuals suggested that discrepant IQ profiles may relate to variations in structural (39) and functional organization (40). These relationships, however, have not been systematically assessed in ASD. To fill this gap, the current study investigated structural and functional network substrates of cognitive imbalances in ASD, capitalizing on a multimodal neuroimaging and connectomics approach. Core to our study was a broad battery of analytical approaches, including correlative analyses with nvIQ ratios, verbal and non-verbal IQ profile clustering, and dimensional IQ profile decomposition. We leveraged the autism brain imaging data exchange initiative datasets (ABIDE-I and II; 41,42), which offer the currently largest repository containing imaging and phenotypic information. Analyses were regionally unconstrained, operating at a cortex- and connectome-wide level; we nevertheless hypothesized that imbalance of verbal and non-verbal dimensions of intelligence would particularly manifest in the structure and functional network embedding of language networks, but also in those involved in sensory and higher cognitive functions. Moreover, we expected that the different data science techniques would overall converge on a robust structure-function substrate of cognitive imbalances in autism.

## Results

We studied two aggregate datasets of individuals with ASD and neurotypical individuals from both waves of the Autism Brain Imaging Data Exchange initiative (ABIDE-I and -II; http://fcon_1000.projects.nitrc.org/indi/abide; 41,42).

*Dataset-1* was used for main phenotypic and neuroimaging analyses. Inclusion criteria were similar to our prior imaging studies in ASD (22,23,36,43,44). Specifically, we focused on those sites that included both children and adults with an autism diagnosis or who were neurotypical controls in sufficient numbers. Excluding cases with low-quality structural MRI or inaccurate cortical surface extraction (visually checked and manually corrected by SLV and BCB) resulted in a final sample of 155 ASD (150 males, mean±SD age in years for ASD =17.9±8.6) and 151 controls (150 males; mean±SD age in years=17.7±7.3). Data was from four different sites (*i.e*., PITT [n=42], USM [n=92], NYU [n=135], TCD [n=37]). Quality indices for structural and functional MRI data did not differ between ASD and controls (p>0.43, t=0.79 for cortical surface extraction, p>0.11, t=1.58 for head motion). Details on subject inclusion and quality control are provided in the *Methods* and **Figure S1**.

*Dataset-2* was an independent selection from the ABIDE-I and II waves, and included individuals that were not included in the imaging sample but that had IQ measures available (325 ASD; mean±SD age in years =15.8±7.3; 284 males; 569 controls, 12.7±5.6 years, 420 males). Phenotypic replication findings from this cohort are presented below as well.

### Analysis of vnIQ ratio data and its structure-function substrate

We first studied the ratio of verbal over non-verbal IQ (vnIQ) to capture cognitive imbalance in a single score. Studying *Dataset-1*, ASD showed a reduced vnIQ ratio compared to controls (*Cohen’s d*=0.27, p<0.016), mainly due to markedly decreased verbal IQ (*d*=0.72, p<0.0001) and less marked changes in non-verbal IQ (*d*=0.45, p<0.0001). The prevalence of imbalanced verbal and non-verbal IQ (*i.e*., verbal IQ < non-verbal IQ) was also higher in ASD relative to neurotypicals (χ^2^=4.67, p<0.03). Findings were largely reproduced in the *Dataset-2* (nvIQ reduction: *d*=0.18, p<0.015; verbal IQ reduction: *d*=0.35, p<0.0001; non-verbal IQ reduction: *d*=0.16, p<0.03; prevalence: χ^2^=8.83, p<0.005).

MRI-based cortical thickness analysis explored morphological substrates of imbalanced cognitive profiles. Specifically, we assessed surface-wide interactions between the diagnostic groups (*i.e*., ASD *vs*. neurotypicals) and vnIQ ratio, and identified significant clusters in the left insular, fronto-opercular, paracentral and posterior midline cortices as well as the right fronto-opercular cortices (**Figure 2A**). In these regions, increased thickness in neurotypicals was associated with increased vnIQ, while ASD showed an inverse pattern with atypical thickening relating to lower scores (*i.e*., more severe cognitive imbalance). Post-hoc analyses focusing on significant clusters confirmed moderate cross-site generalizability of this interaction, showing variable effects yet with a consistent direction across included sites (PITT: t=1.57, p<0.065; USM: t=1.81, p<0.04; NYU: t=2.45, p<0.01; TCD: t=1.25, p<0.12; **FigureS2A**). We observed a similar interaction when splitting our dataset into children and adults, with however slightly more marked effects in children (Children: t=3.77, p<0.001; Adults: t=1.75, p<0.045; **Figure S2B**). Notably, spatial decoding of the map showing significant group by vnIQ interactions via automated meta analyses (http://neurosynth.org; 45) identified multiple terms, with ‘language’ being the top-ranked one, but also lower level perceptual and somato-motor processes (*e.g*., ‘pain’, ‘motor’, ‘auditory processing’) as well as higher order cognitive processes (*e.g*., ‘semantics’, ‘declarative memory’, ‘cognitive control’). Similarly, we leveraged a recently proposed intrinsic functional community parcellation (*i.e*., the Cole-Anticevic atlas that explicitly includes a set of parcels for the language network; (46) and could confirm a marginal interaction between vnIQ and diagnostic group (ASD, controls) on cortical thickness in language networks (t=1.7, p<0.057; **Figure S2C**).

**Figure 1.**
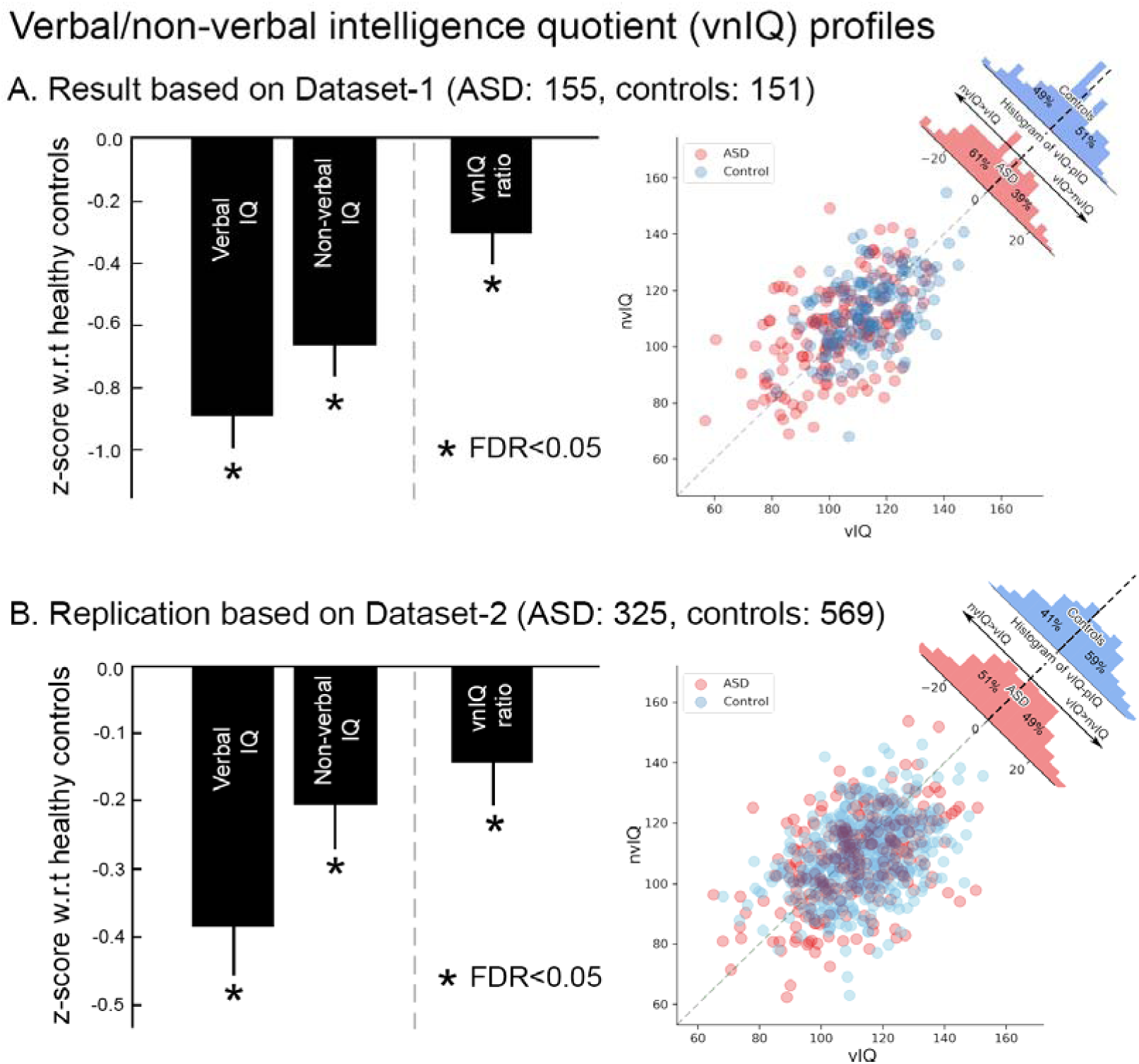
Verbal and non-verbal IQ profiles in ASD. **A) Findings in Dataset-1** *Left*. Z-scores of verbal IQ, nonverbal IQ, and the vnIQ ratio (verbal/nonverbal IQ) in ASD relative to neurotypical controls. Error bars present standard deviations. *Right*. The prevalence of imbalanced verbal and nonverbal IQ (*i.e*., verbal < non-verbal IQ). **B) Phenotypic replication in Dataset-2**.

**Figure 2.**
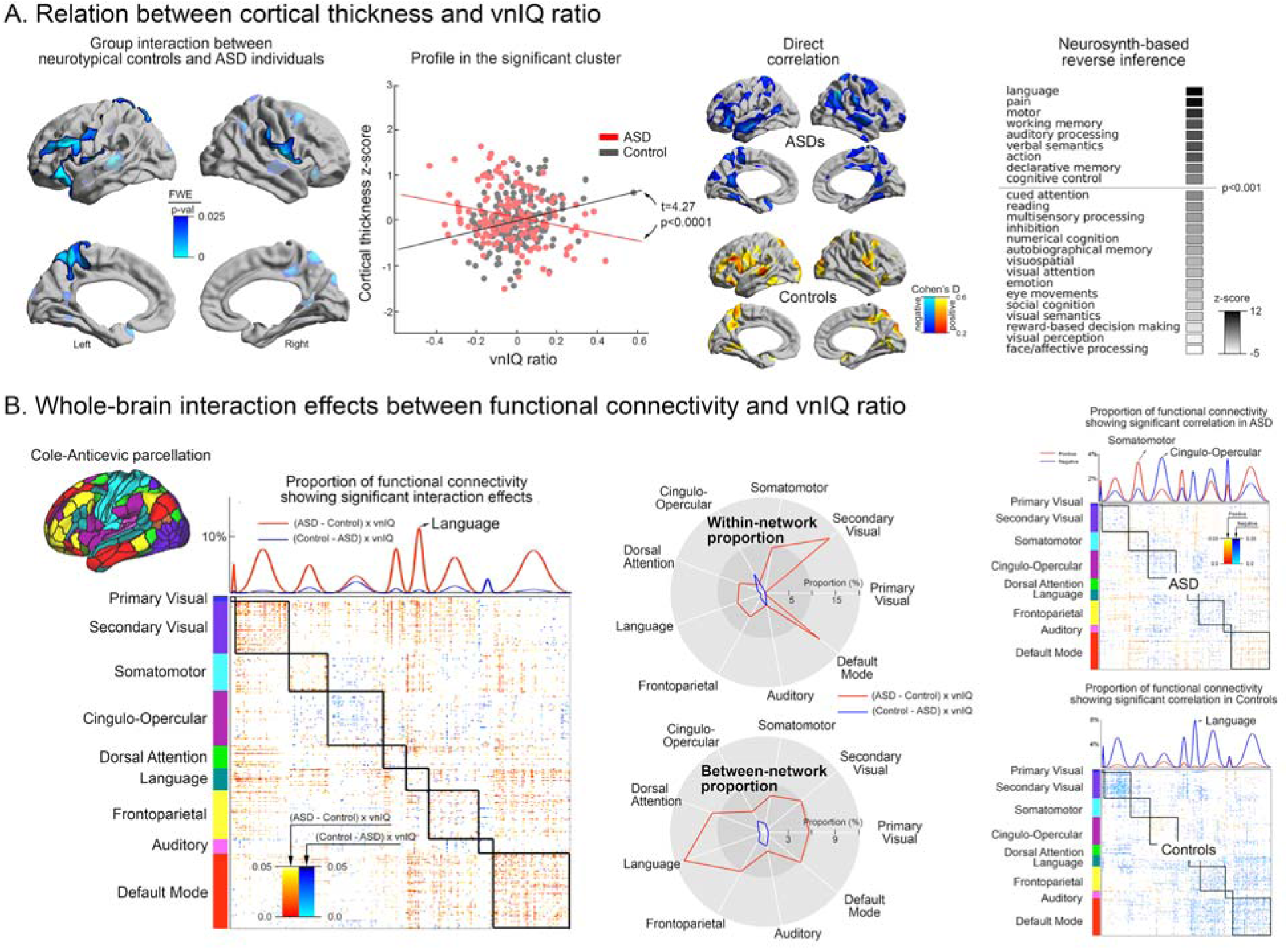
Structure-function substrates of vnIQ imbalance in ASD. **A) Cortical thickness analysis** *Left*. Interaction between diagnostic group (ASD, controls) and vnIQ scores. FWE-corrected clusters are shown in solid and with black outlines, uncorrected trends in semitransparent. *Center left*. Post-hoc analysis on mean thickness in significant clusters. *Center right*. Correlation between cortical thickness and vnIQ in each group separately. *Right*. Significant clusters from the group-by-vnIQ interaction (see *Left*) analysis were fed into Neurosynth and associated with cognitive terms. **B) Functional associations**. *Left*. Using a functional parcellation that includes the language network (46), interaction analysis was performed at the level of parcel-to-parcel connections. Uncorrected p-values from this interaction analysis were sorted according to functional communities (primary visual, secondary visual, somatomotor, cingulo-opercular, dorsal attention, language, frontoparietal, auditory, default mode). Parcel-wise significant interactions were summed within each network, and stratified into within-vs. between-community connections. *Right*. Direct correlation analysis between vnIQ and functional connectivity across different communities, carried out in ASD and controls separately. Positive/negative effects are indicated in blue/red.

Next, we evaluated associations of functional network organization and cognitive imbalances. Connectome-wide resting-state fMRI analysis investigated connectivity profiles within and between different functional communities, using the same intrinsic functional community classification as above (47). We confirmed interactions between vnIQ and diagnostic group (ASD, controls) on the connectivity of multiple networks (**Figure 2B left and middle**). Stratification of effects with respect to within/between community communication highlighted alterations of within-community communication for default mode (mean±SD t=2.12±0.40) and sensory (mean±SD t=2.09±0.34) networks, while particularly the language network (mean±SD t=2.07±0.35) displayed interaction effects in connections to other networks. We also explored connectome-wide modulations in each group separately (**Figure 2B right**). In ASD, the vnIQ ratio modulated positively and negatively the connectivity in multiple networks, while associations in controls were mainly negative. Specifically, in ASD somatomotor, visual, and default mode showed more frequently positive associations, while cingulo-opercular, language, and auditory networks showed rather negative associations. Functional interaction effects were consistent when varying preprocessing choices, specifically when additionally controlling for global mean signal (**Figure S3**). As for cortical thickness findings, effects were relatively consistent across the included sites (**Figure S4A**) and when repeating the analyses within children or adults (**Figure S4B**).

### Categorical subtyping of IQ profiles and structure-function substrates

We complemented the vnIQ interaction analyses with categorical subtyping to identify discrete boundaries across ASD individuals in terms of cognitive imbalances. Agglomerative hierarchical clustering and silhouette analyses based on verbal and non-verbal IQ provided solutions with 2 or 4 clusters as compact and most appropriate. While the two-subtype solution split the individuals with ASD along the overall cognitive performance (*i.e*., low *vs*. high full-scale IQ), the four-subtype solution revealed one additional axis related to cognitive imbalances (reduced *vs*. increased vnIQ; **Figure 3A**). Given its relevance to our hypothesis, we focused on this four-subtype solution. ASD1 was low full-scale IQ but with slightly imbalanced verbal and non-verbal IQ (z-score of the vnIQ ratio relative to neurotypical controls=-0.79), whereas ASD3 was a high (or average) full-scale IQ subgroup without marked cognitive imbalance (z-score of the vnIQ ratio=-0.45). On the other hand, ASD2 and ASD4, demonstrated dichotomized patterns (*i.e*., z-score of the vnIQ ratio in ASD2/ASD4=-1.12/0.85) with less marked change in terms of general cognitive performance (mean±SD full-scale IQ=106±13/107±14). Notably, we observed a similar phenotype subtype solution in the independent Dataset-2 (**Figure S5**).

**Figure 3.**
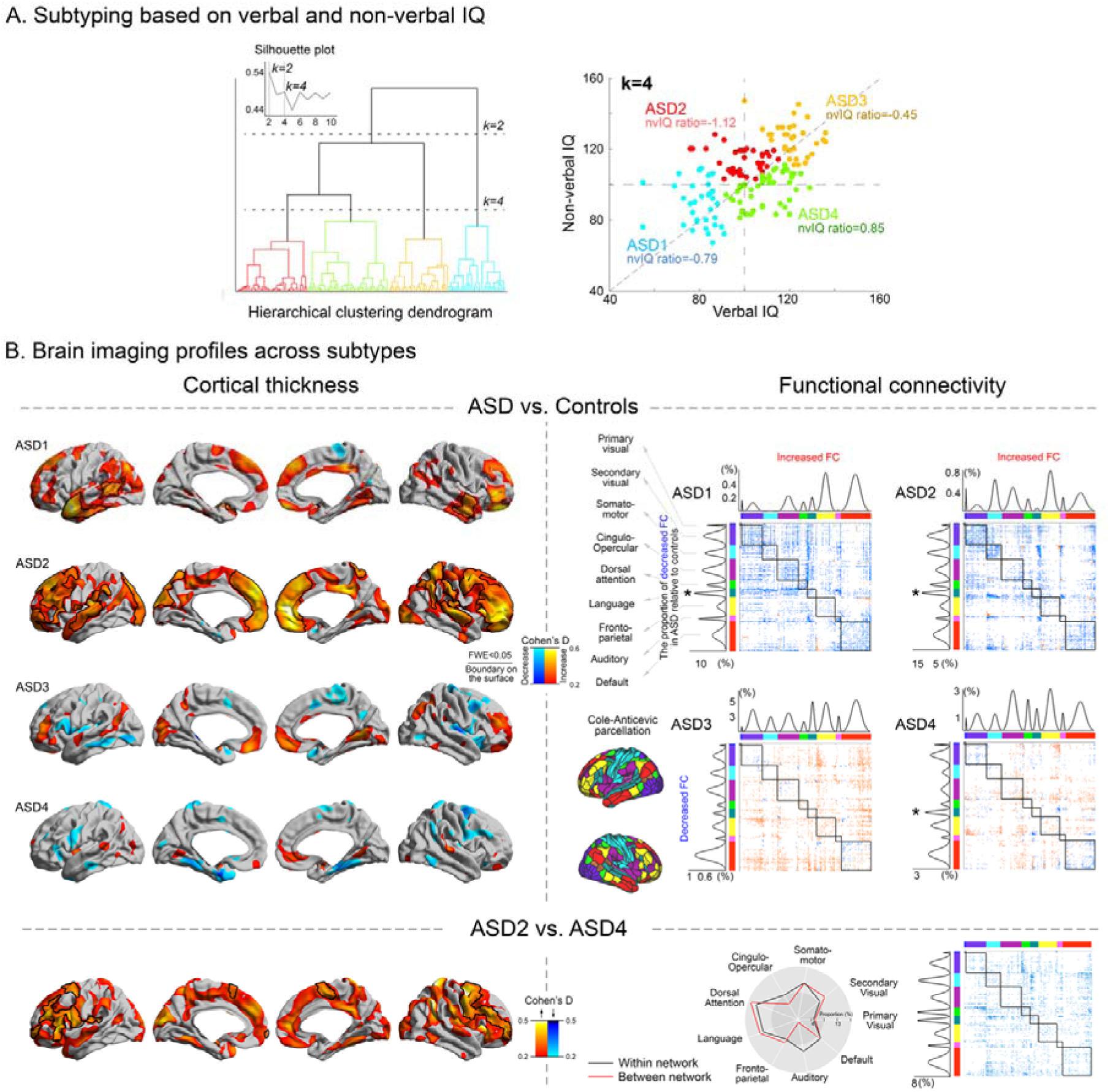
Subtyping based on verbal and non-verbal IQ. **A)** Subtyping results shown at k=2 and 4, the solutions resulting in the highest Silhouette index. **B)** A k=4 solution provided subtypes reflecting cognitive imbalances, particularly when comparing ASD2 vs ASD4. Cortical thickness and functional connectivity features were profiled across all four subtypes, by comparing these measures to neurotypical controls (effect sizes are presented as Cohen’s *D*). For the functional connectivity analysis, connections showing a significant between-group difference were counted within- and between-network separately and plotted left to the connectome for within-community comparisons and above the corresponding connectome for between-community analyses, for each subtype. The peak modulation in the language network is marked by * and was observed in 3 out of the 4 identified subtypes. The bottom panels show targeted comparisons between ASD2 (high non-verbal compared to verbal IQ) and ASD4 (high verbal compared to non-verbal IQ).

The IQ-derived ASD subtypes presented with differential cortical thickness alterations relative to neurotypical controls, in a spectrum that encompassed widespread cortical thickness increases (ASD1 and ASD2) together with more localized patchy patterns of increases and decreases in thickness (ASD3 and ASD4) (**Figure 3B left**). Notably, contrasting ASD2 and ASD4 indicated widespread cortical thickness in ASD2, the class that also showed the lowest vnIQ ratio compared to the other ASD subgroups. Mirroring the findings from the vnIQ analysis, it should be noted that these increases unlikely reflect simple effects of lower verbal IQ. Indeed, when comparing ASD1 (*i.e*., the group with more reduced verbal IQ than ASD2) and ASD4, we did not observe marked thickness increases, neither cortex-wide nor in language networks (**Figure S6**). Considering functional connectivity, all subtypes presented with variable patterns of decreased as well as increased connectivity across multiple networks relative to controls. The classes ASD1 and ASD2 that also showed increased cortical thickening presented with largely functional connectivity reductions relative to controls (mean Cohen’s *D*=0.34/0.33 for within- and between-community connectivity, respectively), while ASD3 and ASD4 showed mixed patterns of both increases (*D*=0.31/0.30 for within- and between-community connectivity, respectively) and decreases (*D*=0.31/0.29 for within-/between-community). Most marked modulations were again observed in language networks (**Figure 3B right**). Directly contrasting ASD2 and ASD4 revealed more marked hypoconnectivity in ASD2 compared to ASD4 (*D*=0.34/0.33 for within-/between-community connectivity).

### Dimensional IQ subtyping and convergent structure-function associations

We finally ran a principal component analysis of verbal and non-verbal IQs in ASD to tap into biological continuity (48), we (**Figure 4**). Two principal components were identified, each of which explained distinct dimensions of ASD IQ profile variance (PC1: 76%, PC2: 24%). Specifically, PC1 reflected more closely the average of verbal to non-verbal IQ while PC2 reflected more closely its imbalance. Notably, PC2 showed a clear spectrum (spanning from ASD2 to ASD4 from the above subtyping analysis), with one extreme characterized by high functioning non-verbal IQ yet together with low verbal IQ as well as the other extreme showing enhanced verbal ability but with reduced non-verbal IQ profiles. We observed similar components in the held-out dataset (**Figure S7**).

**Figure 4.**
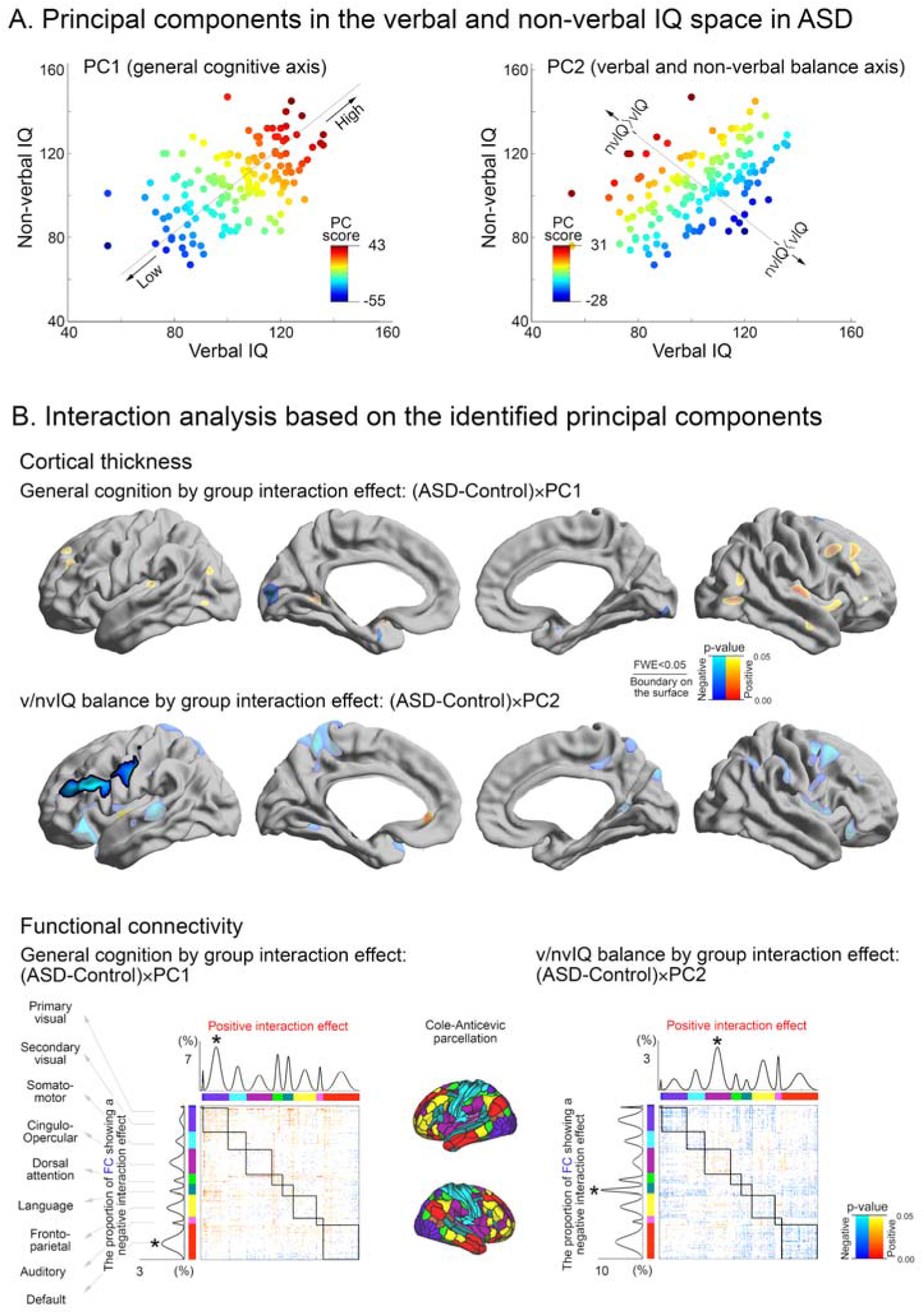
Principal component analysis for the distribution of verbal and non-verbal IQ in ASD. **A)** The direction and scores of the two principal components derived from verbal and non-verbal IQ profiles. PC1 reflected individual variability along a general cognitive axis and PC2 reflected verbal to non-verbal IQ imbalance. **B)** Between-group interaction analysis of PC1 and PC2 on cortical thickness and functional connectivity. Upper panels show significant cortical thickness modulations were delineated by solid boundaries whereas uncorrected tendencies are shown in semi-transparent. Lower panels show the proportion of functional connections that undergo a significant between-group interaction for both PC1 and PC2. Findings were stratified according to functional communities as in the prior Figures.

Group interaction analyses between PC scores and brain imaging features revealed that *i)* in PC1, the ASD group presented atypically more cortical thickening associated with higher general cognitive performance compared to the neurotypical control group, and *ii)* in PC2, the increased thickness was coupled with severe vnIQ imbalance (more deficits in verbal compared to non-verbal IQ) with higher degree in ASD than in neurotypicals. Functional connectivity also revealed similar group-dependent changes with respect to its relationship with each PC score. For the general cognitive axis (PC1), both positive effects were most marked in the visual network (t=1.96/1.97 for within- and between-community connectivity respectively), while negative effects predominated in transmodal systems such as default mode network (t=2.02/1.97 for within- and between-community connectivity respectively). Notably, however, for the PC2 axis (verbal/non-verbal imbalance), communities demonstrating significant interaction effects were either the language network (higher PC2 scores and thus more severe vnIQ reduction, related to decreased connectivity in ASD relative to controls; t=1.92/2.01 for within-/between-community connectivity) or the salience network (higher the PC2 scores related to connectivity increases in ASD relative to controls; t=1.95/1.95 for within-/between-community connectivity), confirming the tight relationship between the observed cognitive imbalance and the language-related brain areas.

## Discussion

Inter-individual heterogeneity in biological, cognitive, and behavioral dimensions is increasingly recognized to hinder research and intervention in autism (3,7,22,49). The current diagnostic classification of ASD is relatively broad, and has been suggested to capture different severities and potential etiologies (2,8). While this approach may overall increase diagnostic sensitivity, it may on the other hand reduce specificity motivating additional stratification to further calibrate diagnostics and guide intervention. Here, we targeted autism heterogeneity using a “*cognitive-first*” perspective by studying verbal and non-verbal dimensions of intelligence in multi-site datasets of individuals with ASD and neurotypical controls. In contrast to the current use of IQ measures as clinical specifiers without diagnostic indication, but in line with emerging observations of markedly discrepant cognitive profiles in autism (5,9,14), the current work provided robust evidence for a marked cognitive imbalance in ASD compared to neurotypicals. Indeed, capitalizing on three alternative data analytical strategies (*i.e*., linear associations to vnIQ ratio, IQ profile clustering, dimensional IQ profile decomposition), we could show in a large multicentric cohort that individuals with ASD presented with markedly imbalanced verbal to non-verbal intelligence. Moreover, harnessing state-of-the-art brain imaging and connectomics, we could demonstrate that imbalances converged on a structure-function substrate characterized by atypical morphology and functional network embedding of language, sensory-motor, and higher cognitive systems. Phenotypic findings could be replicated in an independent cohort, suggesting generalizability. Our findings, thus, provide robust evidence that verbal and non-verbal intelligence disparities are markedly accentuated in ASD. Moreover, they outline a neural substrate of this imbalance in the connectivity and morphology of the language system alongside with multiple lower as well as higher order networks. These findings overall motivate the incorporation of cognitive imbalances in autism research, which may capture a broad range of autism phenotypes and potentially help to stratify individuals prior to interventions.

Our first series of analyses quantified cognitive imbalances by calculating a simple ratio between verbal and non-verbal IQ profiles, an approach previously adopted in phenotype analyses of typical and atypical development (10,15). Studying two independent cohorts aggregated from the ABIDE-I and II waves, we could find robust evidence for a marked imbalance in ASD. While imbalances were present in both directions, the highest discrepancy was driven by markedly lower verbal compared to non-verbal IQ. Studying the modulations of vnIQ profiles on structural and functional neuroimaging measures, we observed a marked discrepancy between ASD and controls. At the level of MRI-based cortical thickness, we observed marked between-group interactions in regions involved in language networks *i.e*., an inverse relation between vnIQ and thickness in ASD and controls. Specifically, while neurotypical individuals with a high vnIQ ratio presented with higher thickness in language networks, a negative association was observed in ASD, where decreased vnIQ scores reflected atypical thickening. This finding also extended to individuals for those with normal non-verbal intelligence, indicating that this does not reflect the effect of overall intellectual disability. Meta-analytical decoding using neurosynth revealed that the spatial distribution of these findings was indeed aligned to networks involved in language processing. This finding is in line with the intention of the vnIQ ratio measure to be sensitive to language related impairments in the presence of normal non-verbal functioning that has been frequently described in ASD (50). Structural MRI analysis was complemented with an rs-fMRI paradigm to assess functional network substrates of ASD-related cognitive imbalances. Resting-state fMRI has been proposed as a candidate to supply intermediate phenotypes of complex neurodevelopmental conditions such as ASD (26,51–53), as it allows the assessment of whole-brain connectivity at the level of individual subjects. As for the structural MRI findings, this independent approach also pointed to a broad cortical substrate, showing marked interactions between diagnostic group and vnIQ on intrinsic functional organization. Indeed, connectome-wide analysis revealed that functional connections were differentially modulated by vnIQ in ASD relative to controls. These effects were found both within and between multiple macroscopic functional communities showing for example modulations of within-community connectivity within sensory/motor and transmodal networks such as the default mode network. Yet, and in convergence with the structural MRI findings, the analysis also pointed to highest effects in the language network, particularly at the level of its connectivity to other communities.

To dispel potential analysis-related variations, we furthermore typified IQ imbalances using two complementary data analytics, namely clustering-based subtyping and dimensional decomposition of verbal and non-verbal IQ values. Clustering is increasingly applied to identify categorical subgroups in heterogeneous conditions such as autism, epilepsy, and depression (22,54–56). In ASD, an emerging body of studies has attempted to subtype cohorts using behavioral and neuroimaging measures (3,8,22,57,58), including work from our group that clustered a multi-site cohort of individuals with ASD and neurotypicals based on structural MRI markers of horizontal and vertical cortical organization (22). Categorical approaches have been complemented by dimensional assessments, previously proposed as an overarching neuropsychiatric research strategy (59). Here, individual variations are mapped across continuous axes instead of delineating discrete boundaries, between subtypes. Despite their analytical differences, both techniques confirmed modulatory effects of cognitive imbalances in ASD on structural and functional network organization. Indeed, although IQ subtypes were also found to reflect overall cognitive functioning (for example, when considering the two-subtype solution or when comparing ASD-1 and ASD-3 in the four-subtype solution), a more granular four-subtype solution highlighted cognitive imbalances specifically in ASD-2 vs ASD-4, the two subtypes with relatively normal full-scale IQ. Yet, the ASD-2 showed reduced verbal functioning, while ASD-4 presented with the opposite pattern. Direct comparison of both subtypes was in line with the above vnIQ interaction findings, showing fronto-central cortical thickening in individuals with lower verbal IQ, together with connectivity anomalies related to between-network connectivity in these individuals. These findings were corroborated by a subsequent principal components analysis (PCA) on individual IQ profiles. In contrast to clustering, PCA identifies continuous dimensions contributing to IQ profile variability in ASD. In our data, we identified a principal dimension scaling with overall IQ, as well as an orthogonal second dimension sensitive to cognitive imbalances. This latter component reflects brain structure and function in a similar manner as the vnIQ interaction and clustering findings, pointing to cortical thickening and functional connectivity modulations of lateral frontal language networks by degrees of cognitive imbalance in ASD. By showing virtually identical structural and functional connectivity findings across these three analytical strategies, our study thus provides robust evidence of a structure-function substrate of cognitive imbalances in ASD.

Our neuroimaging findings suggest that cognitive imbalance may represent an important source of phenotypic heterogeneity in autism, and that its non-consideration might have contributed to inconclusive findings across previous case-control studies in ASD. Indeed, while some prior structural MRI findings suggested altered thickening in frontal and temporal cortices in ASD relative to neurotypicals (23,25,32,60), other studies have shown cortical thinning or only subtle effects (33), overall leading to a limited consistency (61). Similar to the structural imaging literature, many prior rs-fMRI work followed a case-control design without incorporation of IQ profiles (62–64). Specifically, some reports emphasized reductions in cortico-cortical and long-range connections (36,65,66), while some others suggested atypical organization of local connectivity patterns (29) and in subcortical-cortical connectivity (44,67–69). While such a divergence can partially be attributed to methodological choices and motion-related related confounds on connectivity findings (30,70), a contribution may stem from an inherent heterogeneity in the composition of the ASD cohort (3,7,22,57,71,72,73,74). A recent model that consolidates behavioral, neuroimaging, and genetic findings suggests that genetic mutations may trigger brain reorganization in individuals with a low plasticity threshold, such as transmodal association cortices that have a high number of synapses (16). These reorganizational patterns, and their role in the cortical hierarchies may furthermore lead to alterations in perceptual processing pathways. Such a cascading effect may account for alterations in high-level cognitive processes, ranging from enhancements to impairments (75), changes in visual and auditory perceptual processing (76,77), and for difficulties in more integrative, social cognitive functions in autism. It may thus be tempting to speculate that findings showing differential modulations of brain structure and function by cognitive imbalance in ASD may be underpinned by a developmental mechanism that is similar to cross-modal compensation. However, as neurobiological substrates of cortical thickness and functional connectivity anomalies remain relatively unclear in ASD at this point, it remains to be investigated how far our findings reflect ASD-related effects on brain network plasticity. Early post-mortem work suggested intracortical cellular and laminar anomalies (78,79), together with ectopic neurons in the white matter (80,81). These findings are complemented by work showing alterations in dendritic spine densities on cortical projection neurons in ASD, showing increased dendritic spine densities in supragranular layers in frontal regions and in infragranular layers in temporal cortices (82). These authors discussed the possibility that increased spine densities may result from deficient postnatal culling of connections and may thus have downstream effects on the interplay of excitation and inhibition in local microcircuits, which may be in line with recent connectome modelling studies (44,83). Other reports have shown focal ‘patches’ of disorganized cortical layers in ASD (84) and emphasized altered columnar arrangement, again with prominent findings in temporal and frontal cortices (85,86).

We close by highlighting that our study does not imply that researchers need to control for IQ measures as a variable of no interest when comparing individuals with ASD to neurotypical controls. As our results emphasize, verbal and non-verbal aspects of intelligence and their disparities are instead an important dimension of the diverse autism phenotype, which encompasses a high prevalence of impaired as well as enhanced abilities compared to neurotypicals (16). Although etiological factors that contribute to these imbalances remain to be further investigated, our findings showing robust structure function-substrate of imbalances point to atypical large-scale brain reorganization centered around language related networks, potentially downstream to perturbations of cross-network plasticity.

## Methods and materials

### Participants

*Dataset-1*: We studied a subsample of individuals with ASD and neurotypical individuals from the first two waves of the Autism Brain Imaging Data Exchange initiative (ABIDE-I and -II; 41,42). Inclusion criteria were similar to our prior studies (22,23,36,43,44); specifically, we restricted our assessment to those sites that included both children and adults, with ≥10 individuals per diagnostic group (n=406, 203/203 ASD/controls). Data availability and detailed quality control criteria were then used to select only the cases with verbal/performance IQ scores and acceptable MRI quality (see *below*). Moreover, we additionally removed the data from one site (Institut Pasteur) as the site contained only ASD without matched controls. This series of screening process resulted in 306 individuals (155 ASD [age mean±SD=17.9±8.6, 150 males], 151 controls [age=17.7±7.3, 150 males]) from four different sites: 1) NYU Langone Medical Center (NYU, 65/70 ASD/controls); 2) University of Utah, School of Medicine (USM, 52/40 ASD/controls); 3) University of Pittsburgh, School of Medicine (PITT, 20/22 AS/controls); 4) Trinity Centre for Health Sciences, Trinity College Dublin (TCD, 18/19 ASD/controls).

*Dataset-2*: From all other remaining sites, we aggregated verbal and non-verbal IQ scores among those participants who were not included in the main analyses (n=894, 325/569 ASD/controls).

Individuals with ASD underwent structured or unstructured in-person interview and had a diagnosis of Autistic, Asperger’s, or Pervasive Developmental Disorder Not-Otherwise-Specified established by expert clinical opinion aided by ‘gold standard’ diagnostics: Autism Diagnostic Observation Schedule, ADOS and/or Autism Diagnostic Interview-Revised, ADI-R. These focus on three domains including reciprocal social interactions, communication and language, and restricted/repeated behaviors and interests. Full scale/non-verbal IQ/verbal IQ (105±16 vs. 115±13/106±17 vs. 113±13/103±17 vs. 114±13 for ASD vs. controls, respectively) was measured via WASI, WAIS III, and/or WISC III. Controls had no history of mental disorders and were statistically matched for age to the ASD group at each site. In the ABIDE-I and –II datasets, there were no differences in age and sex between controls and ASD. ABIDE-I and -II datasets are based on studies approved by local IRBs, and data were fully anonymized (removing all HIPAA protected health information identifiers, and face information from structural images).

### MRI acquisition

High-resolution T1-weighted images (T1w) and resting-state functional MRIs (rs-fMRI) were available from all sites. Additionally, the sites from ABIDE-II (*i.e*., NYU, TCD) also included diffusion weighted images (DWI). Images were acquired on 3T scanners from Siemens (NYU, USM, PITT) or Philips (IP, TCD). *NYU* data were acquired on an Allegra using 3D-TurboFLASH for T1w (TR=2530ms; TE=3.25ms; TI=1100ms; flip angle=7**°**; matrix=256×256; 1.3×1.0×1.3mm^3^ voxels), 2D-EPI for rs-fMRI (TR=2000ms; TE=15ms; flip angle=90**°**; matrix=80×80; 180 volumes, 3.0×3.0×4.0mm^3^ voxels) and SE-EPI for DWI (TR=5200ms; TE=78ms; flip angle=60°; axial slices=50; slice thickness=3 mm; voxels=3.0×3.0×3.0 mm^3^; directions=64; b0=1000s/mm^2^). *PITT* data were acquired on an Allegra using 3D-MPRAGE for T1w (TR=2100ms; TE=3.93ms; TI=1000ms; flip angle=7**°**; matrix=269×269; 1.1×1.1×1.1mm^3^ voxels) and 2D-EPI for rs-fMRI (TR=1500ms; TE=35ms; flip angle=70**°**; matrix=64×64; 200 volumes, 3.1×3.1×4.0mm^3^ voxels). *USM* data were acquired on a TrioTim using 3D-MPRAGE for T1w (TR=2300ms; TE=2.91ms; TI=900ms; flip angle=9**°**; matrix=240×256; 1.0×1.0×1.2mm^3^ voxels) and 2D-EPI for rs-fMRI (TR=2000ms; TE=28ms; flip angle=90**°**; matrix=64×64; 240 volumes; 3.4×3.4×3.0mm^3^ voxels). *TCD* data were acquired on an Achieva using 3D-MPRAGE for T1w (TR=3000ms; TE=3.90ms; TI=1150ms; flip angle=8**°**; matrix=256×256; 0.9×0.9×0.9mm^3^ voxels), 2D-EPI for rs-fMRI (TR=2000ms; TE=27ms; flip angle=90**°**; matrix=80×80; 210 volumes; 3.0×3.0×3.2mm^3^ voxels) and SE-EPI for DWI (TR=20244ms; TE=79ms; matrix=124×124; FOV=248; slice thickness=2mm; flip angle=90°; voxels=1.94×1.94×2.0mm^3^; directions=61; b0=1500s/mm^2^).

### MRI processing

This work capitalized on established multimodal image processing and co-registration routines to analyze structural, functional, and diffusion MRI data in the same reference frame.

a. *Structural MRI*. Processing of T1w images was based on FreeSurfer (v5.1; http://surfer.nmr.mgh.harvard.edu/; 87). Image processing included bias field correction, registration to stereotaxic space, intensity normalization, skull-stripping, and white matter segmentation. A triangular surface tessellation fitted a deformable mesh onto the white matter volume, providing grey-white and pial surfaces with >160,000 corresponding vertices. We measured cortical thickness as the distance between grey-white and pial surfaces and registered individual surfaces to template surface, fsaverage5, improving correspondence of measurements with respect to sulco-gyral patterns. Thickness data were smoothed using a surface-based kernel with a full-width-at-half-maximum (FWHM) of 20mm, as in prior studies (88–90).
b. *rs-fMRI*. We leveraged data disseminated via the Preprocessed Connectomes initiative (http://preprocessed-connectomes-project.org/abide/). Processing was based on the Configurable Pipeline for the Analysis of Connectomes, CPAC (https://fcp-indi.github.io/), and included slice-time correction, head motion correction, skull stripping, and intensity normalization. Statistical corrections removed effects of head motion, white matter and cerebro-spinal fluid signals (using the CompCor tool, based on the top 5 principal components; 91), as well as linear/quadratic trends. After band-pass filtering (0.01-0.1Hz), we co-registered rs-fMRI and T1w data through combined linear and non-linear transformations. Surface alignment was verified for each case and we interpolated voxel-wise rs-fMRI time-series along the mid-thickness surface model. We resampled rs-fMRI surface data to Conte69, a template mesh from the Human Connectome Project pipeline (https://github.com/Washington-University/Pipelines) and applied a 5mm FWHM surface-based smoothing.

Multimodal processing was followed by detailed quality control routines. In brief, all subjects with severe faulty surface segmentations, or imaging artifacts, or more than 0.3mm framewise displacement in the functional scans were excluded. Minor segmentation inaccuracies of all remaining cases were manually corrected by several raters (SV, BB), and pipelines rerun.

### Dimensional assessment of brain substrates underlying cognitive profiles

We first assessed differences in verbal and non-verbal IQ as well as their ratio (vnIQ=verbal IQ divided by non-verbal IQ) in ASD relative to controls, controlling for age and site effects. A series of analyses then assessed the effects of vnIQ on cortical morphology and connectivity using the SurfStat toolbox (http://www.math.mcgill.ca/~keith/surfstat (92) for Matlab (The Mathworks, Natick). To then evaluate morphological substrates of IQ profiles, we assessed correlations between the vnIQ ratio in both typically developing controls and ASD, and assessed group-by-vnIQ ratio interactions. Age and site effects were statistically controlled in these models. Ad-hoc meta-analyses identified cognitive term associations of the identified interaction effects, using neurosynth.org (45). To further establish functional associations, we also conducted following two analyses targeting *i)* connectivity modulation effects by vnIQ ratio (*i.e*., regression analysis), and *ii)* critically the interactions between vnIQ ratio and diagnostic group on functional connectivity across the whole brain, as identical to the previous structural MRI analysis. As before, we controlled for the effects of age and site. In keeping with our recent functional connectivity work in ASD (36), we included framewise displacement in the linear model. All surface-based anatomical findings were corrected for multiple comparisons using random field theory for non-isotropic images (93).

### Categorical approach: data-driven clustering analysis based on the IQ ratio and subtype profiling

The previous analyses assessed a group-common relationship between the IQ ratio and brain imaging features based on entire ASD and control subjects. Given an increasingly more adapted perspective on substantial biological heterogeneity in autism, however, our study also evaluated whether the IQ ratio reveals detectable discrete boundaries across individuals with ASD. To this purpose, we carried out a categorical subtyping approach of IQ measures by applying agglomerative hierarchical clustering analysis (kernel: *Wald’s linkage*) on verbal IQ, non-verbal IQ, and vnIQ. Silhouette analysis benchmarked the clustering solution. Identified subtypes were then comprehensively profiled in terms of both IQ scores and brain imaging features. Especially for the latter, we have compared the cortical thickness as well as whole-brain functional connectivity between ASD individuals in each subtype and neurotypical controls to see if cognitive-driven subtypes reveal distinct patterns in underlying system-level neurobiology. Notably, our post-hoc analysis selected two subtypes that best recapitulate our main interests of hypothesis (*i.e*., a subtype showing the lowest vnIQ and the other showing the highest vnIQ) and directly compared their multimodal imaging features to visualize the effect of cognitive imbalance in ASD individuals with more increased sensitivity.

### Dimensional approach: principal component analysis based on the IQ ratio and imaging profiles

Another perspective more recently adapted in the field to better understand the autistic condition is a dimensional approach, which emphasizes biological continuity on its pathological mechanism rather than assuming that ASD consists of clearly separable multiple subtypes. To assess the utility of this complementing approach, we applied a principal component analysis (PCA) on the two targeted IQ measures (verbal and non-verbal IQ) and mapped the component scores to the individuals to sort them out along the identified IQ axes. In other words, the component score assigned to each individual serves as an indicator where the individual is ranked in the v-and nv-IQ space along the identified principal axis. Given the number of input features (=2; verbal IQ and non-verbal IQ) to PCA, the total components identifiable were also two. As done in the previous subtyping analyses, we related individual component scores to brain imaging features by performing a group interaction analysis based on a linear model (*i.e*., [ASD-control or control-ASD]×PC-scores) in both cortical thickness and functional connectivity.

## Acknowledgments

This work was supported by funding from the Brain & Behavior Research Foundation (NARSAD Young Investigator grant; #28436), the Canadian Institutes of Health Research (postdoctoral fellowship MFE-158228) and the Institute for Basic Science (IBS-R15-D1) in Korea for SJH. SL acknowledges funding from Fonds de la Recherche du Québec – Santé (FRQ-S) and the Canadian Institutes of Health Research (CIHR). RVdW receives support from a Savoy Foundation studentship. BP is supported by a Molson Engineering Fellowship and the National Research Foundation of Korea (NRF-2020R1A6A3A03037088). IS acknowledges funding from FRQ-S. AM acknowledges research funding from NIEHS (R01 ES030950, R01 ES027424, K23ES026239), NIH (1UG3OD023290), and the NVLD Project. BCB acknowledges research funding from the SickKids Foundation (NI17-039), the National Sciences and Engineering Research Council of Canada (NSERC; Discovery-1304413), CIHR (FDN-154298), Azrieli Center for Autism Research (ACAR), an MNI-Cambridge collaboration grant, BrainCanada (Azrieli Future Leaders), FRQS, and the Canada Research Chairs Tier 2 program.

## References

1. Maenner MJ, Shaw KA, Baio J, EdS1, Washington A, Patrick M, et al. (2020): Prevalence of Autism Spectrum Disorder Among Children Aged 8 Years - Autism and Developmental Disabilities Monitoring Network, 11 Sites, United States, 2016. MMWR Surveill Summ 69: 1–12.

2. Mottron L, Bzdok D (2020): Autism spectrum heterogeneity: fact or artifact? Mol Psychiatry. https://doi.org/10.1038/s41380-020-0748-y

3. Lombardo MV, Lai M-C, Baron-Cohen S (2019): Big data approaches to decomposing heterogeneity across the autism spectrum. Mol Psychiatry 24: 1435–1450.

4. Hong SJ, Vogelstein JT, Gozzi A, Bernhardt BC (2020): Towards Neurosubtypes in Autism. Biologicals. Retrieved from https://www.sciencedirect.com/science/article/pii/S0006322320314979

5. Mottron L, Dawson M, Soulières I, Hubert B, Burack J (2006): Enhanced perceptual functioning in autism: an update, and eight principles of autistic perception. J Autism Dev Disord 36: 27–43.

6. Baron-Cohen S (2017): Editorial Perspective: Neurodiversity - a revolutionary concept for autism and psychiatry. J Child Psychol Psychiatry 58: 744–747.

7. Rødgaard E-M, Jensen K, Vergnes J-N, Soulières I, Mottron L (2019): Temporal Changes in Effect Sizes of Studies Comparing Individuals With and Without Autism: A Meta-analysis. JAMA Psychiatry. https://doi.org/10.1001/jamapsychiatry.2019.1956

8. Lai M-C, Lombardo MV, Chakrabarti B, Baron-Cohen S (2013): Subgrouping the autism “spectrum”: reflections on DSM-5. PLoS Biol 11: e1001544.

9. Munson J, Dawson G, Sterling L, Beauchaine T, Zhou A, Elizabeth K, et al. (2008): Evidence for latent classes of IQ in young children with autism spectrum disorder. Am J Ment Retard 113: 439–452.

10. Ankenman K, Elgin J, Sullivan K, Vincent L, Bernier R (2014): Nonverbal and verbal cognitive discrepancy profiles in autism spectrum disorders: influence of age and gender. Am J Intellect Dev Disabil 119: 84–99.

11. Coolican J, Bryson SE, Zwaigenbaum L (2008): Brief Report: Data on the Stanford–Binet Intelligence Scales (5th ed.) in Children with Autism Spectrum Disorder. Journal of Autism and Developmental Disorders, vol. 38. pp 190–197.

12. Joseph RM, Tager-Flusberg H, Lord C (2002): Cognitive profiles and social-communicative functioning in children with autism spectrum disorder. J Child Psychol Psychiatry 43: 807–821.

13. Nowell KP, Brewton CM, Allain E, Mire SS (2015): The Influence of Demographic Factors on the Identification of Autism Spectrum Disorder: A Review and Call for Research. Review Journal of Autism and Developmental Disorders, vol. 2. pp 300–309.

14. Ostrolenk A, Forgeot d’Arc B, Jelenic P, Samson F, Mottron L (2017): Hyperlexia: Systematic review, neurocognitive modelling, and outcome. Neurosci Biobehav Rev 79: 134–149.

15. Nader A-M, Jelenic P, Soulières I (2015): Discrepancy between WISC-III and WISC-IV Cognitive Profile in Autism Spectrum: What Does It Reveal about Autistic Cognition? PLoS One 10: e0144645.

16. Mottron L, Belleville S, Rouleau GA, Collignon O (2014): Linking neocortical, cognitive, and genetic variability in autism with alterations of brain plasticity: the Trigger-Threshold-Target model. Neurosci Biobehav Rev 47: 735–752.

17. Silbereis JC, Pochareddy S, Zhu Y, Li M, Sestan N (2016): The Cellular and Molecular Landscapes of the Developing Human Central Nervous System. Neuron 89: 248–268.

18. Lariviere S, Vos de Wael R, Paquola C, Hong S-J, Mišić B, Bernasconi N, et al. (2019): Microstructure-informed connectomics: enriching large-scale descriptions of healthy and diseased brains. Brain Connect 9: 113–127.

19. Betzel RF, Bassett DS (2017): Multi-scale brain networks. Neuroimage 160: 73–83.

20. Gilmore JH, Knickmeyer RC, Gao W (2018): Imaging structural and functional brain development in early childhood. Nature Reviews Neuroscience, vol. 19. pp 123–137.

21. Lerch JP, van der Kouwe AJW, Raznahan A, Paus T, Johansen-Berg H, Miller KL, et al. (2017): Studying neuroanatomy using MRI. Nat Neurosci 20: 314–326.

22. Hong S-J, Valk SL, Di Martino A, Milham MP, Bernhardt BC (2018): Multidimensional Neuroanatomical Subtyping of Autism Spectrum Disorder. Cereb Cortex 28: 3578–3588.

23. Valk SL, Di Martino A, Milham MP, Bernhardt BC (2015): Multicenter mapping of structural network alterations in autism. Hum Brain Mapp 36: 2364–2373.

24. Raznahan A, Toro R, Daly E, Robertson D, Murphy C, Deeley Q, et al. (2010): Cortical anatomy in autism spectrum disorder: an in vivo MRI study on the effect of age. Cereb Cortex 20: 1332–1340.

25. Bedford SA, Park MTM, Devenyi GA, Tullo S, Germann J, Patel R, et al. (2020): Large-scale analyses of the relationship between sex, age and intelligence quotient heterogeneity and cortical morphometry in autism spectrum disorder. Mol Psychiatry 25: 614–628.

26. Di Martino A, Fair DA, Kelly C, Satterthwaite TD, Castellanos FX, Thomason ME, et al. (2014): Unraveling the miswired connectome: a developmental perspective. Neuron 83: 1335–1353.

27. Craddock RC, Jbabdi S, Yan C-G, Vogelstein JT, Castellanos FX, Di Martino A, et al. (2013): Imaging human connectomes at the macroscale. Nat Methods 10: 524–539.

28. Di Martino A, Kelly C, Grzadzinski R, Zuo X-N, Mennes M, Mairena MA, et al. (2011): Aberrant striatal functional connectivity in children with autism. Biol Psychiatry 69: 847–856.

29. Keown CL, Shih P, Nair A, Peterson N, Mulvey ME, Müller R-A (2013): Local functional overconnectivity in posterior brain regions is associated with symptom severity in autism spectrum disorders. Cell Rep 5: 567–572.

30. Müller R-A, Shih P, Keehn B, Deyoe JR, Leyden KM, Shukla DK (2011): Underconnected, but how? A survey of functional connectivity MRI studies in autism spectrum disorders. Cereb Cortex 21: 2233–2243.

31. van Rooij D, Anagnostou E, Arango C, Auzias G, Behrmann M, Busatto GF, et al. (2018): Cortical and Subcortical Brain Morphometry Differences Between Patients With Autism Spectrum Disorder and Healthy Individuals Across the Lifespan: Results From the ENIGMA ASD Working Group. Am J Psychiatry 175: 359–369.

32. Hardan AY, Muddasani S, Vemulapalli M, Keshavan MS, Minshew NJ (2006): An MRI study of increased cortical thickness in autism. Am J Psychiatry 163: 1290–1292.

33. Wallace GL, Dankner N, Kenworthy L, Giedd JN, Martin A (2010): Age-related temporal and parietal cortical thinning in autism spectrum disorders. Brain 133: 3745–3754.

34. Haar S, Berman S, Behrmann M, Dinstein I (2014): Anatomical Abnormalities in Autism? Cereb Cortex 26: 1440–1452.

35. Holiga Š, Hipp JF, Chatham CH, Garces P, Spooren W, D’Ardhuy XL, et al. (2019): Patients with autism spectrum disorders display reproducible functional connectivity alterations. Sci Transl Med 11. https://doi.org/10.1126/scitranslmed.aat9223

36. Hong S-J, Vos de Wael R, Bethlehem RAI, Lariviere S, Paquola C, Valk SL, et al. (2019): Atypical functional connectome hierarchy in autism. Nat Commun 10: 1022.

37. Uddin LQ, Supekar K, Menon V (2013): Reconceptualizing functional brain connectivity in autism from a developmental perspective. Front Hum Neurosci 7: 458.

38. King JB, Prigge MBD, King CK, Morgan J, Weathersby F, Fox JC, et al. (2019): Generalizability and reproducibility of functional connectivity in autism. Mol Autism 10: 27.

39. Margolis A, Bansal R, Hao X, Algermissen M, Erickson C, Klahr KW, et al. (2013): Using IQ discrepancy scores to examine the neural correlates of specific cognitive abilities. J Neurosci 33: 14135–14145.

40. Margolis AE, Davis KS, Pao LS, Lewis A, Yang X, Tau G, et al. (2018): Verbal-spatial IQ discrepancies impact brain activation associated with the resolution of cognitive conflict in children and adolescents. Dev Sci 21. https://doi.org/10.1111/desc.12550

41. Di Martino A, Yan C-G, Li Q, Denio E, Castellanos FX, Alaerts K, et al. (2014): The autism brain imaging data exchange: towards a large-scale evaluation of the intrinsic brain architecture in autism. Mol Psychiatry 19: 659–667.

42. Di Martino A, O’Connor D, Chen B, Alaerts K, Anderson JS, Assaf M, et al. (2017): Enhancing studies of the connectome in autism using the autism brain imaging data exchange II. Sci Data 4: 170010.

43. Hong S-J, Hyung B, Paquola C, Bernhardt BC (2019): The Superficial White Matter in Autism and Its Role in Connectivity Anomalies and Symptom Severity. Cereb Cortex 29: 4415–4425.

44. Park B-Y, Hong S-J, Valk S, Paquola C, Benkarim O, Bethlehem RAI, et al. (n.d.): Connectome and microcircuit models implicate atypical subcortico-cortical interactions in autism pathophysiology. https://doi.org/10.1101/2020.05.08.077289

45. Yarkoni T, Poldrack RA, Nichols TE, Van Essen DC, Wager TD (2011): Large-scale automated synthesis of human functional neuroimaging data. Nat Methods 8: 665.

46. Ji JL, Spronk M, Kulkarni K, Repovš G, Anticevic A, Cole MW (2019): Mapping the human brain’s cortical-subcortical functional network organization. Neuroimage 185: 35–57.

47. Ji JL, Spronk M, Kulkarni K, Repovš G, Anticevic A, Cole MW (2019): Mapping the human brain’s cortical-subcortical functional network organization. Neuroimage 185: 35–57.

48. Sestan N, State MW (2018): Lost in Translation: Traversing the Complex Path from Genomics to Therapeutics in Autism Spectrum Disorder. Neuron 100: 406–423.

49. Hong S-J, Vogelstein JT, Gozzi A, Bernhardt BC, Thomas Yeo BT, Milham MP, Di Martino A (2020): Toward Neurosubtypes in Autism. Biological Psychiatry, vol. 88. pp 111–128.

50. Dawson M, Soulières I, Gernsbacher MA, Mottron L (2007): The level and nature of autistic intelligence. Psychol Sci 18: 657–662.

51. Castellanos FX, Xavier Castellanos F, Di Martino A, Cameron Craddock R, Mehta AD, Milham MP (2013): Clinical applications of the functional connectome. NeuroImage, vol. 80. pp 527–540.

52. Parkes L, Satterthwaite TD, Bassett DS (2020): Towards precise resting-state fMRI biomarkers in psychiatry: synthesizing developments in transdiagnostic research, dimensional models of psychopathology, and normative neurodevelopment. Curr Opin Neurobiol 65: 120–128.

53. Uddin LQ, Supekar K, Menon V (2010): Typical and atypical development of functional human brain networks: insights from resting-state FMRI. Front Syst Neurosci 4: 21.

54. Drysdale AT, Grosenick L, Downar J, Dunlop K, Mansouri F, Meng Y, et al. (2017): Resting-state connectivity biomarkers define neurophysiological subtypes of depression. Nat Med 23: 28–38.

55. Bernhardt BC, Hong S-J, Bernasconi A, Bernasconi N (2015): Magnetic resonance imaging pattern learning in temporal lobe epilepsy: classification and prognostics. Ann Neurol 77: 436–446.

56. Hong S-J, Lee H-M, Gill R, Crane J, Sziklas V, Bernhardt BC, et al. (2019): A connectome-based mechanistic model of focal cortical dysplasia. Brain 142: 688–699.

57. Hong S-J, Vogelstein JT, Gozzi A, Bernhardt BC, Thomas Yeo BT, Milham M, di Martino A (n.d.): Towards Neurosubtypes in Autism. https://doi.org/10.31234/osf.io/8az69

58. Feczko E, Balba NM, Miranda-Dominguez O, Cordova M, Karalunas SL, Irwin L, et al. (2018): Subtyping cognitive profiles in Autism Spectrum Disorder using a Functional Random Forest algorithm. Neuroimage 172: 674–688.

59. Insel T, Cuthbert B, Garvey M, Heinssen R, Pine DS, Quinn K, et al. (2010): Research domain criteria (RDoC): toward a new classification framework for research on mental disorders. Am J Psychiatry 167. https://doi.org/10.1176/appi.ajp.2010.09091379

60. van Rooij D, Anagnostou E, Arango C, Auzias G, Behrmann M, Busatto GF, et al. (2018): Cortical and Subcortical Brain Morphometry Differences Between Patients With Autism Spectrum Disorder and Healthy Individuals Across the Lifespan: Results From the ENIGMA ASD Working Group. Am J Psychiatry 175: 359–369.

61. Duerden EG, Mak-Fan KM, Taylor MJ, Roberts SW (2012): Regional differences in grey and white matter in children and adults with autism spectrum disorders: an activation likelihood estimate (ALE) meta-analysis. Autism Res 5: 49–66.

62. Cherkassky VL, Kana RK, Keller TA, Just MA (2006): Functional connectivity in a baseline resting-state network in autism. NeuroReport, vol. 17. pp 1687–1690.

63. Monk CS, Peltier SJ, Wiggins JL, Weng S-J, Carrasco M, Risi S, Lord C (2009): Abnormalities of intrinsic functional connectivity in autism spectrum disorders. NeuroImage, vol. 47. pp 764–772.

64. Washington SD, Gordon EM, Brar J, Warburton S, Sawyer AT, Wolfe A, et al. (2014): Dysmaturation of the default mode network in autism. Hum Brain Mapp 35: 1284–1296.

65. Tomasi D, Volkow ND (2019): Reduced Local and Increased Long-Range Functional Connectivity of the Thalamus in Autism Spectrum Disorder. Cereb Cortex 29: 573–585.

66. Khan S, Gramfort A, Shetty NR, Kitzbichler MG, Ganesan S, Moran JM, et al. (2013): Local and long-range functional connectivity is reduced in concert in autism spectrum disorders. Proc Natl Acad Sci U S A 110: 3107–3112.

67. Cerliani L, Mennes M, Thomas RM, Di Martino A, Thioux M, Keysers C (2015): Increased Functional Connectivity Between Subcortical and Cortical Resting-State Networks in Autism Spectrum Disorder. JAMA Psychiatry 72: 767–777.

68. Nair A, Treiber JM, Shukla DK, Shih P, Müller R-A (2013): Impaired thalamocortical connectivity in autism spectrum disorder: a study of functional and anatomical connectivity. Brain 136: 1942–1955.

69. Mizuno A, Villalobos ME, Davies MM, Dahl BC, Müller R-A (2006): Partially enhanced thalamocortical functional connectivity in autism. Brain Res 1104: 160–174.

70. Nair A, Keown CL, Datko M, Shih P, Keehn B, Müller R-A (2014): Impact of methodological variables on functional connectivity findings in autism spectrum disorders. Hum Brain Mapp 35: 4035–4048.

71. Benkarim O, Paquola C, Park B-Y, Hong S-J, Royer J, de Wael RV, et al. (n.d.): Functional idiosyncrasy has a shared topography with group-level connectivity alterations in autism. https://doi.org/10.1101/2020.12.18.423291

72. Bernhardt BC, Di Martino A, Valk SL, Wallace GL (2017): Neuroimaging-Based Phenotyping of the Autism Spectrum. Curr Top Behav Neurosci 30: 341–355.

73. Nunes AS, Peatfield N, Vakorin V, Doesburg SM (2019): Idiosyncratic organization of cortical networks in autism spectrum disorder. Neuroimage 190: 182–190.

74. Dickie EW, Ameis SH, Shahab S, Calarco N, Smith DE, Miranda D, et al. (2018): Personalized Intrinsic Network Topography Mapping and Functional Connectivity Deficits in Autism Spectrum Disorder. Biol Psychiatry 84: 278–286.

75. Mottron L, Bouvet L, Bonnel A, Samson F, Burack JA, Dawson M, Heaton P (2013): Veridical mapping in the development of exceptional autistic abilities. Neurosci Biobehav Rev 37: 209–228.

76. Wang L, Mottron L, Peng D, Berthiaume C, Dawson M (2007): Local bias and local-to-global interference without global deficit: a robust finding in autism under various conditions of attention, exposure time, and visual angle. Cogn Neuropsychol 24: 550–574.

77. O’Connor K (2011): Auditory Processing in Autism Spectrum Disorder: A Review of the Literature: A Thesis Submitted in Partial Fulfilment of the Requirements for the Degree of Master of Audiology at the Department of Communication Disorders, University of Canterbury.

78. Hutsler JJ, Love T, Zhang H (2007): Histological and Magnetic Resonance Imaging Assessment of Cortical Layering and Thickness in Autism Spectrum Disorders. Biological Psychiatry, vol. 61. pp 449–457.

79. Bauman M, Kemper TL (1985): Histoanatomic observations of the brain in early infantile autism. Neurology 35: 866–874.

80. Bauman ML, Kemper TL (2008): The Neuropathology of the Autism Spectrum Disorders: What Have We Learned? Autism: Neural Basis and Treatment Possibilities. pp 112–128.

81. Avino TA, Hutsler JJ (2010): Abnormal cell patterning at the cortical gray-white matter boundary in autism spectrum disorders. Brain Res 1360: 138–146.

82. Hutsler JJ, Zhang H (2010): Increased dendritic spine densities on cortical projection neurons in autism spectrum disorders. Brain Res 1309: 83–94.

83. Trakoshis S, Martínez-Cañada P, Rocchi F, Canella C, You W, Chakrabarti B, et al. (2020): Intrinsic excitation-inhibition imbalance affects medial prefrontal cortex differently in autistic men versus women. Elife 9. https://doi.org/10.7554/eLife.55684

84. Stoner R, Chow ML, Boyle MP, Sunkin SM, Mouton PR, Roy S, et al. (2014): Patches of disorganization in the neocortex of children with autism. N Engl J Med 370: 1209–1219.

85. Casanova MF, Buxhoeveden DP, Switala AE, Roy E (2002): Minicolumnar pathology in autism. Neurology 58: 428–432.

86. Casanova MF, van Kooten IAJ, Switala AE, van Engeland H, Heinsen H, Steinbusch HWM, et al. (2006): Minicolumnar abnormalities in autism. Acta Neuropathol 112: 287–303.

87. Fischl B (2012): FreeSurfer. NeuroImage, vol. 62. pp 774–781.

88. Lerch JP, Evans AC (2005): Cortical thickness analysis examined through power analysis and a population simulation. Neuroimage 24: 163–173.

89. Valk SL, Bernhardt BC, Trautwein F-M, Böckler A, Kanske P, Guizard N, et al. (2017): Structural plasticity of the social brain: Differential change after socio-affective and cognitive mental training. Sci Adv 3: e1700489.

90. Valk SL, Bernhardt BC, Böckler A, Trautwein F-M, Kanske P, Singer T (2017): Socio-Cognitive Phenotypes Differentially Modulate Large-Scale Structural Covariance Networks. Cereb Cortex 27: 1358–1368.

91. Behzadi Y, Restom K, Liau J, Liu TT (2007): A component based noise correction method (CompCor) for BOLD and perfusion based fMRI. NeuroImage, vol. 37. pp 90–101.

92. Worsley KJ, Taylor JE, Carbonell F, Chung MK, Duerden E, Bernhardt B, et al. (2009): SurfStat: A Matlab toolbox for the statistical analysis of univariate and multivariate surface and volumetric data using linear mixed effects models and random field theory. NeuroImage, vol. 47. p S102.

93. Worsley KJ, Andermann M, Koulis T, MacDonald D, Evans AC (1999): Detecting changes in nonisotropic images. Hum Brain Mapp 8: 98–101.

