## Supplementary Materials for "A convergent structure-function substrate of cognitive imbalances in autism"

*1) Multimodal Imaging and Connectome Analysis Laboratory, McConnell Brain Imaging Centre, Montreal Neurological Institute and Hospital, McGill University, Montreal, Canada; 2) Center for the Developing Brain, Child Mind Institute, New York, NY, USA; 3) Center for Neuroscience Imaging Research, Institute for Basic Science, Sungkyunkwan University, Suwon, South Korea; 4) Department of Biomedical Engineering, Sungkyunkwan University, Suwon, South Korea; 5) Centre de recherche du CIUSSSNIM and Department of Psychiatry, Université de Montréal, Montreal, QC, Canada; 6) Institute of Neuroscience and Medicine (INM-7: Brain and Behaviour), Research Centre Jülich, Germany; 7) Institute of Systems Neuroscience, Heinrich Heine University Düsseldorf, Germany; 8) Department of Psychology, Université du Québec à Montréal, Montréal, QC, Canada;9) Department of Psychiatry, The New York State Psychiatric Institute and the College of Physicians Surgeons, Columbia University, New York, NY, USA; 10) Center for Biomedical Imaging and Neuromodulation, Nathan Kline Institute, NY, USA; 11) Autism Center, Child Mind Institute, New York, NY, USA*

### Supplementary Materials

**Supplementary Table 1.** Clinical and demographic profiles of identified subtypes in the imaging cohort.

| | Subtypes | | | | p-value (ANOVA/Chi$^2$) |
|---|---|---|---|---|---|
| | S1 (n=33) | S2 (n=41) | S3 (n=34) | S4 (n=47) | |
| Age (y)† | 17.8±8.8 | 18.9±7.9 | 19.7±10.7 | 15.8±6.9 | 0.17 |
| Sex (f/m) | 2/31 | 2/39 | 1/34 | 1/46 | 0.23 |
| Site (# of cases) PITT/USM/NYU/TCD | 1/17/13/2 | 6/15/17/3 | 8/8/12/6 | 5/12/23/7 | 0.33 |
| Full scale IQ† | 82±7 | 106±6 | 127±6 | 104±9 | >0.0001 |
| Verbal IQ† | 80±8 | 99±9 | 121±8 | 109±9 | >0.0001 |
| Non-verbal IQ† | 88±12 | 113±7 | 126±9 | 97±9 | >0.0001 |
| vnIQ (z-score)† | -0.8±1.4 | -1.1±0.8 | -0.5±0.8 | 0.9±0.9 | >0.0001 |
| Calibrated ADOS† | 7.2±2.0 | 6.5±2.1 | 6.0±2.0 | 6.1±1.9 | 0.039 |

†: Mean±SD

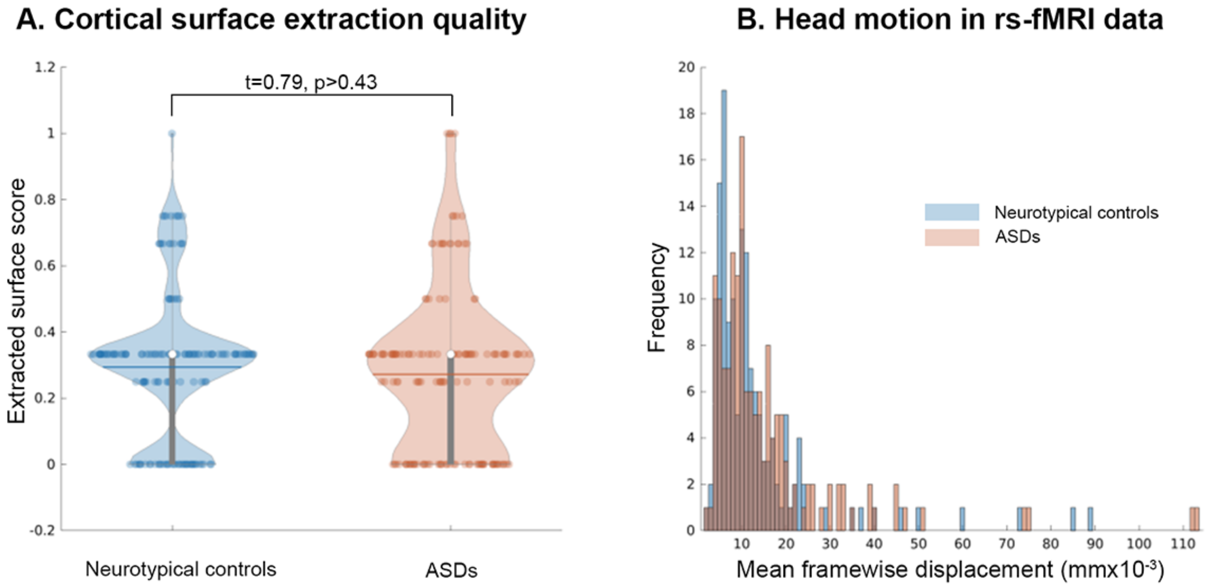


**Supplementary Figure 1. Quality of structural and functional MRI data.**

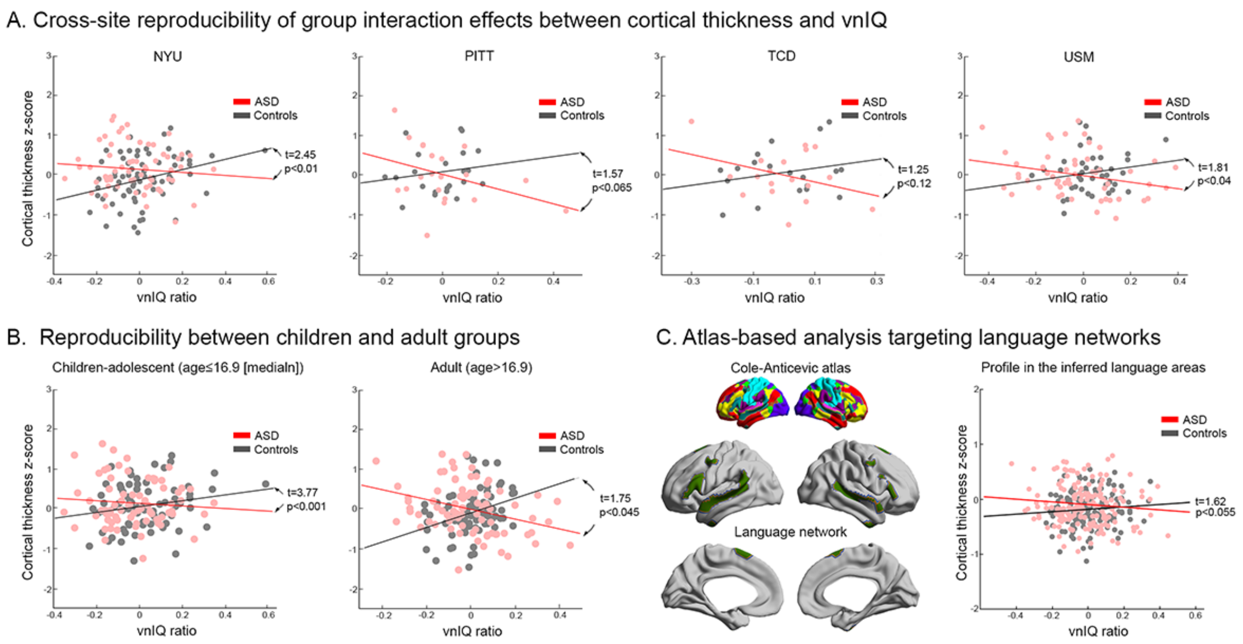


**Supplementary Figure 2. Cross-site and -age reproducibility test for correlative association of cortical thickness to vnIQ ratio**

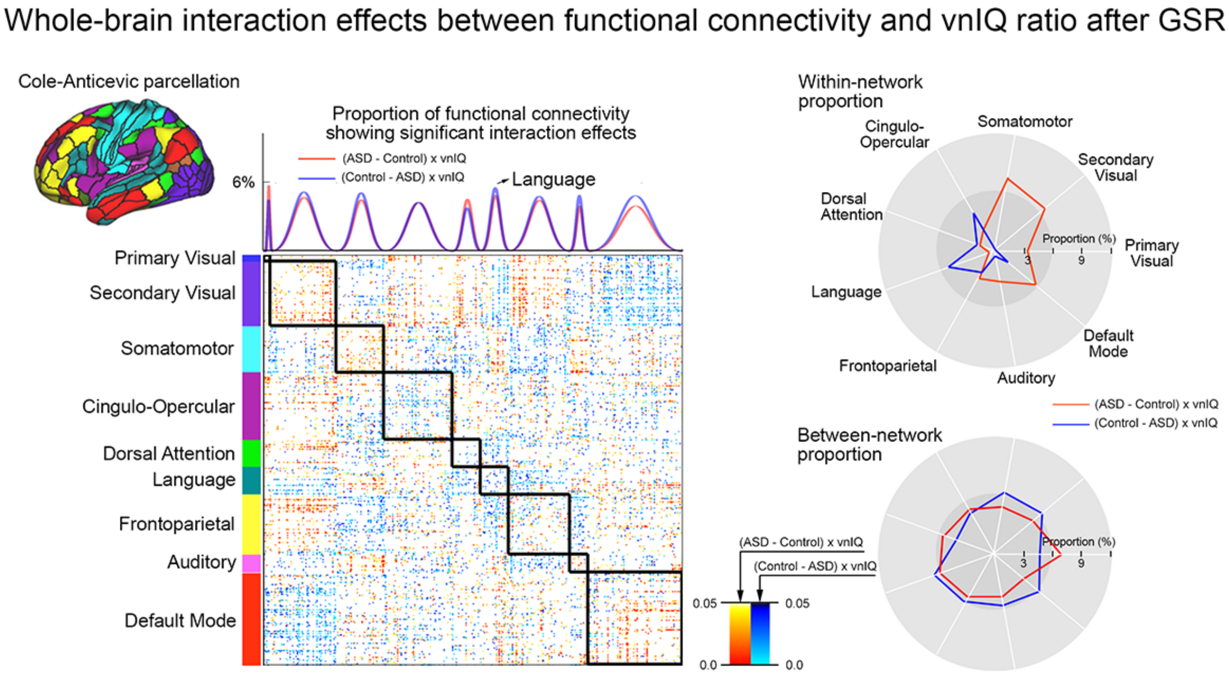


**Supplementary Figure 3. Reproducibility of interaction effect on whole-brain functional connectivity after global mean signal regression**

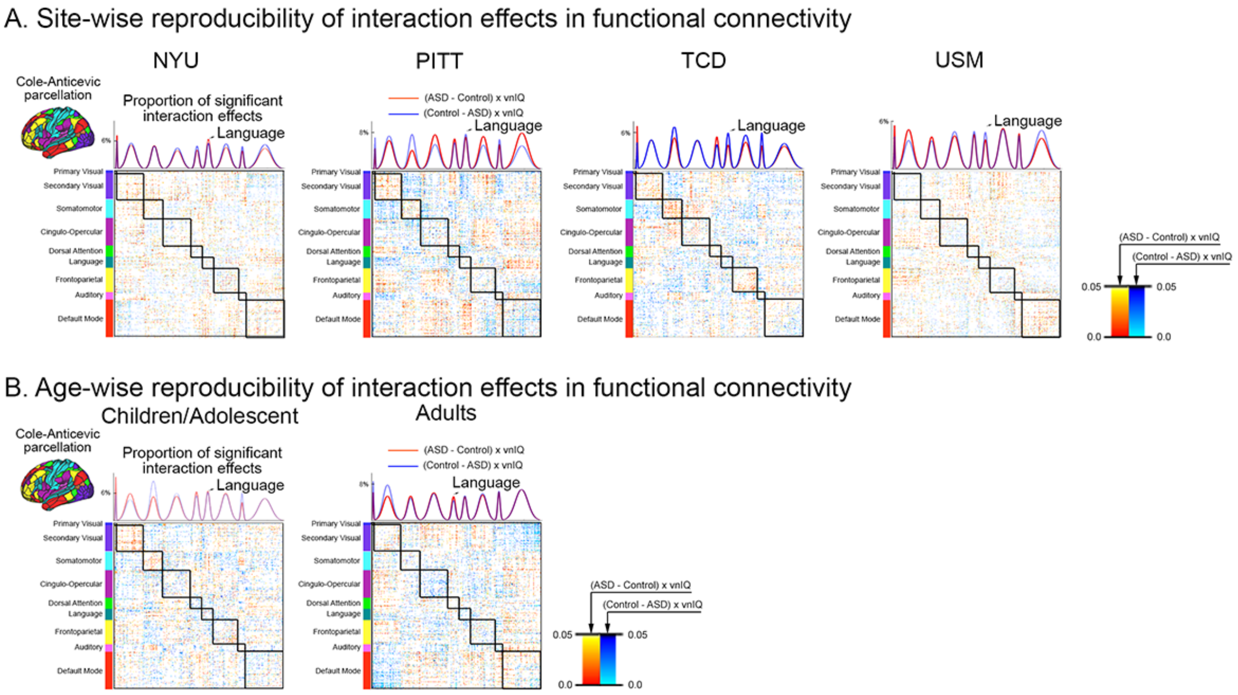


**Supplementary Figure 4. Reproducibility of interaction effect on whole-brain functional connectivity across different sites and age ranges.**

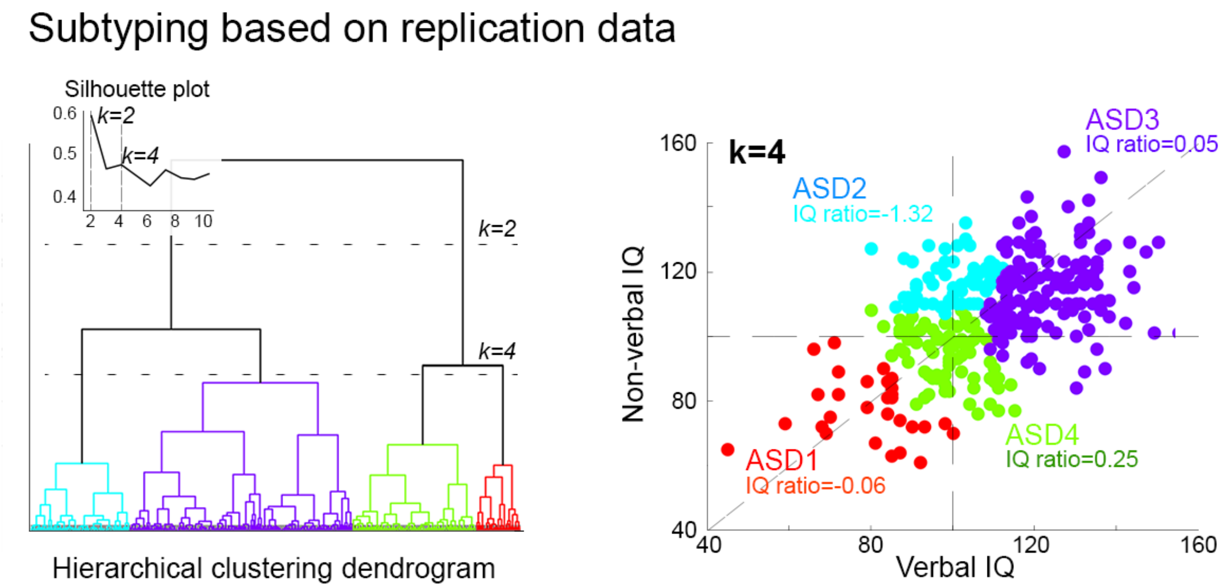


**Supplementary Figure 5. IQ clustering in the independent dataset.**

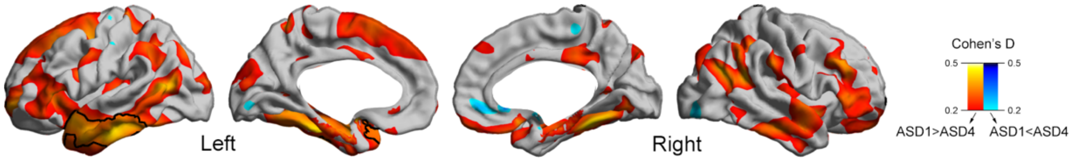


**Supplementary Figure 6. Cortical thickness group comparison between ASD-1 and -4 subtypes.**

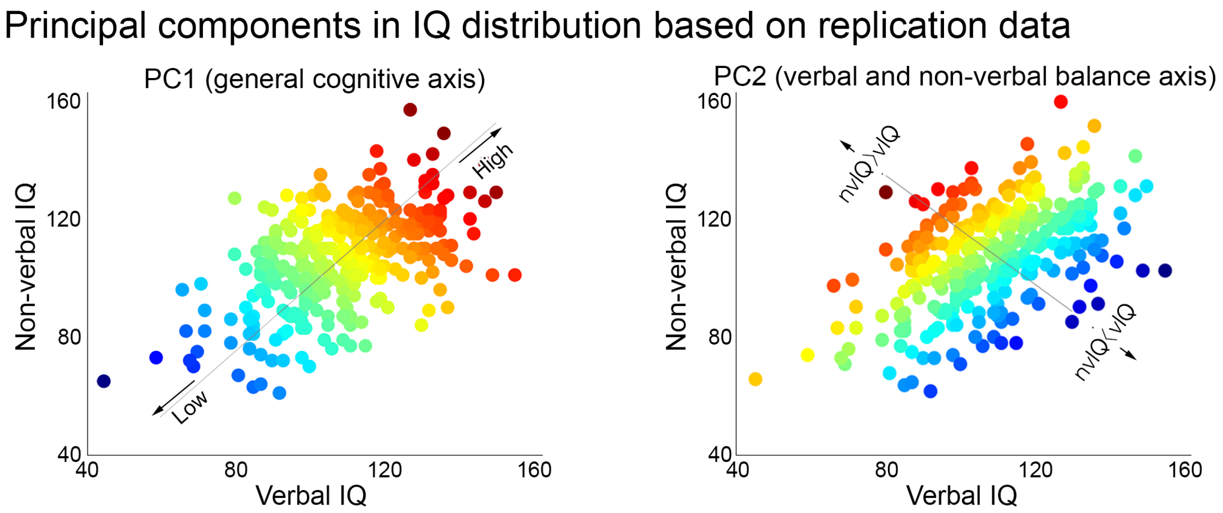


**Supplementary Figure 7. Dimensional IQ components in the independent dataset.**
